# Natural diversity of heat-induced transcription of retrotransposons in *Arabidopsis thaliana*

**DOI:** 10.1101/2024.01.15.575637

**Authors:** Wenbo Xu, Michael Thieme, Anne C. Roulin

## Abstract

Transposable elements (TEs) are major components of plant genomes, profoundly impacting the fitness of their hosts. However, technical bottlenecks have long hindered our mechanistic understanding of TEs. Using RNA-Seq and long-read sequencing with Oxford Nanopore Technologies’ direct cDNA sequencing, we analyzed the heat-induced transcription of TEs in three natural accessions of *Arabidopsis thaliana* (Cvi-0, Col-0, and Ler-1). In addition to the well-studied *ONSEN* retrotransposon family, we identified *Copia-35* as a second heat-responsive retrotransposon family with particularly high activity in the relict accession Cvi-0. Our analysis revealed distinct expression patterns of individual TE copies and suggest different mechanisms regulating the GAG protein production in the *ONSEN* versus *Copia-35* families. In addition, analogously to *ONSEN*, *Copia-35* activation led to the upregulation of flanking genes such as *AMUP9* and potentially to the quantitative modulation of flowering time. Unexpectedly, our results indicate that for both families, the upregulation of flanking genes is not directly initiated by transcription from their 3’ LTRs. These findings highlight the inter- and intraspecific expressional diversity linked to retrotransposon activation under stress, providing insights into their potential roles in plant adaptation and evolution at elevated temperatures.

## Introduction

Transposable elements (TEs) have a profound impact on genome architectures of plants. In crops such as maize, wheat, and barley, TEs account for a majority of the genome, ranging from 64% to more than 80% (Jiao et al. 2017; Wicker et al. 2017; Wicker et al. 2018). Due to their potentially deleterious effects, most TEs are silenced by DNA methylation and through packaging into a heterochromatin state. In particular, one of the most studied plant-specific TE silencing mechanisms is the RNA-directed DNA methylation (RdDM) pathway (Matzke and Mosher 2014). The canonical RdDM pathway features two plant-specific RNA polymerases (Pol IV and Pol V), which, via complex processes, facilitate DNA methylation and, ultimately, the silencing of TEs. Despite widespread silencing, some TEs are still able to transpose in the wild, hereby creating genetic diversity among populations of a given species. For example, a recent study identified ∼23,000 TE insertion polymorphisms (TIPs) across 1047 natural accessions (Baduel et al. 2021) in *Arabidopsis thaliana,* in which TEs account for ∼21% of the genome (Berardini et al., 2015).

Abiotic as well as biotic stresses can provide the conditions that allow specific TE families to evade the host’s silencing mechanisms (Negi et al. 2016). One of the best characterized stress-responsive plant TEs is the retrotransposon (RT) *ONSEN* (or *ATCOPIA78*) in *A. thaliana* (Pecinka et al. 2010; Tittel-Elmer et al. 2010; Ito et al. 2011; Ito et al. 2013). *ONSEN* contains identical long terminal repeats (LTRs) on both ends, as well as coding sequences for gag, the reverse transcriptase and other enzymes, which are essential for its transposition process (Wicker et al. 2007). When *A. thaliana* seedlings are treated with heat, *ONSEN* becomes transcriptionally active, and, upon loss of major epigenetic regulators (Ito et al. 2011) or a transient chemical demethylation (Thieme et al. 2017), it transposes at high frequency, resulting in the stable inheritance of novel *ONSEN* copies.

A particularly interesting feature of *ONSEN* is the fact that its insertions can also confer neighboring genes with heat responsiveness (Ito et al. 2011; Baduel et al. 2021; Roquis et al. 2021), leading to a reshuffling of transcriptional networks. The heat-induced transcription of *ONSEN* flanking genes is attributed to heat-responsive elements in *ONSEN*’s LTRs. These elements recruit heat shock factors that engage the transcription machinery as trimers, resulting in an upregulation of downstream genes (Wu 1995; Cavrak et al. 2014). The finding that *ONSEN* can mediate the expression of flanking regions under heat stress has evolutionary implications since numerous studies have confirmed insertion polymorphisms of *ONSEN* among natural populations (Cavrak et al. 2014; Masuda et al. 2016; Quadrana et al. 2016; Baduel et al. 2021) as well as an insertion bias towards exons and H2A.Z enriched regions (Quadrana et al. 2019; Roquis et al. 2021).

Since the initial discovery of *ONSEN* (Ito et al. 2011), additional heat-responsive TEs have been identified in *A. thaliana*. Two comprehensive experiments using RNA-Seq revealed that in the Col-0 ecotype, both *ONSEN* and *ROMANIAT5* (referred to as *Copia-35* in Repbase) (Pietzenuk et al. 2016; Sun et al. 2020) display heat-dependent transcription. However, while *ONSEN* has been studied in detail, our understanding of *Copia-35* remains limited. A few studies have focused on a particular copy of *Copia-35*, *AT1TE43225*, owing to its role in modulating the expression of its 3’ flanking gene *APUM9*, which encodes the RNA-binding protein *Arabidopsis* PUMILIO9 that triggers the decay of target mRNA (Sanchez and Paszkowski 2014; Hristova et al. 2015). However, the natural diversity of the *APUM9* locus, and more specifically the role of *Copia-35* in driving its expression under heat stress, have not been examined across multiple natural accessions, meaning that our current understanding of the TE contribution to heat-responsiveness is superficial at best.

While technical bottlenecks have been largely responsible for this knowledge gap, the advent of next-generation sequencing now allows to decipher the natural genetic diversity linked to TEs. The availability of polished genome assemblies, produced by long-read sequencing, provides access to the complete sequences of insertions, thereby facilitating a more comprehensive analysis of the genetic features of these insertions. In terms of characterizing the effects of TEs, RNA-Seq has allowed us to survey the entire transcriptome at once, irrespective of the limitations to perceptible phenotypic traits. Technical hurdles persist, however, as the task of aligning short reads from RNA-Seq to multi-copy TEs remains challenging (Lanciano and Cristofari 2020), particularly when the TE copies exhibit a high degree of identity. As a result, transcriptional studies of TEs using RNA-Seq are either based on consensus sequences such as SalmonTE (Jeong et al. 2018) or distribute reads evenly to all copies (Jin et al. 2015). In this context, the breakthrough recently brought by Oxford Nanopore Technologies’ (ONT) direct cDNA sequencing, which generates longer reads, has begun to drastically reduce alignment ambiguities, hereby facilitating the detection of TE expression at the single insertion level. As such, ONT has recently succeeded in improving existing TE annotations. For example, ONT’s cDNA sequencing on a *A. thaliana* mutant with transcriptionally reactivated TEs has allowed to identify and annotate the active TE loci (Panda & Slotkin, 2020). Similarly, long reads generated by ONT recently enabled the identification of chimeric gene-transposon transcripts in *A. thaliana* (Berthelier et al. 2023), further highlighting the advantage of this powerful sequencing technique.

In this study, we examined the patterns of TE expression among natural accessions of heat-stressed *A. thaliana* (particularly between individual TE insertions) and the subsequent effects of TE activation on neighboring genes, by combining the powers of RNA-Seq and Oxford Nanopore Technologies’ (ONT) direct cDNA sequencing. For this purpose, we chose a relict (Cvi-0), a nonrelict (Col-0) and an admixture (Ler-1) accession (Alonso-Blanco et al. 2016). Importantly, each of these accessions previously had polished chromosomal-level PacBio assemblies and annotated genes. Using ONT direct cDNA, we were also able to precisely profile the transcription of heat-activated TEs for the first time in plants. As such, our work not only elucidates the fundamental mechanisms of the stress-induced transcription of TEs but also helps understanding their role as a source of transcriptional novelty and important drivers of evolution.

## Results

### Global comparison of ONT and RNA-seq datasets

We grew Col-0, Ler-1 and Cvi-0 plants under controlled or heat stress conditions and performed RNA-sequencing with classical illumina short-read RNAseq and ONT. We first assessed the data quality of our RNA- and ONT-seq runs (Supplementary Table S1). To verify the effectiveness of the heat stress treatment, we performed a Principal Coordinate Analysis (PCoA) analysis on gene expression using all samples. We found a clear separation of samples based on their treatment and genotype (Fig. 1a), indicating that the applied heat stress induced an accession-specific stress response. Most importantly, this showed that our ONT data was reproducible, and that differences between sequencing technologies did not overshadow global gene expression estimates.

**Figure 1.**
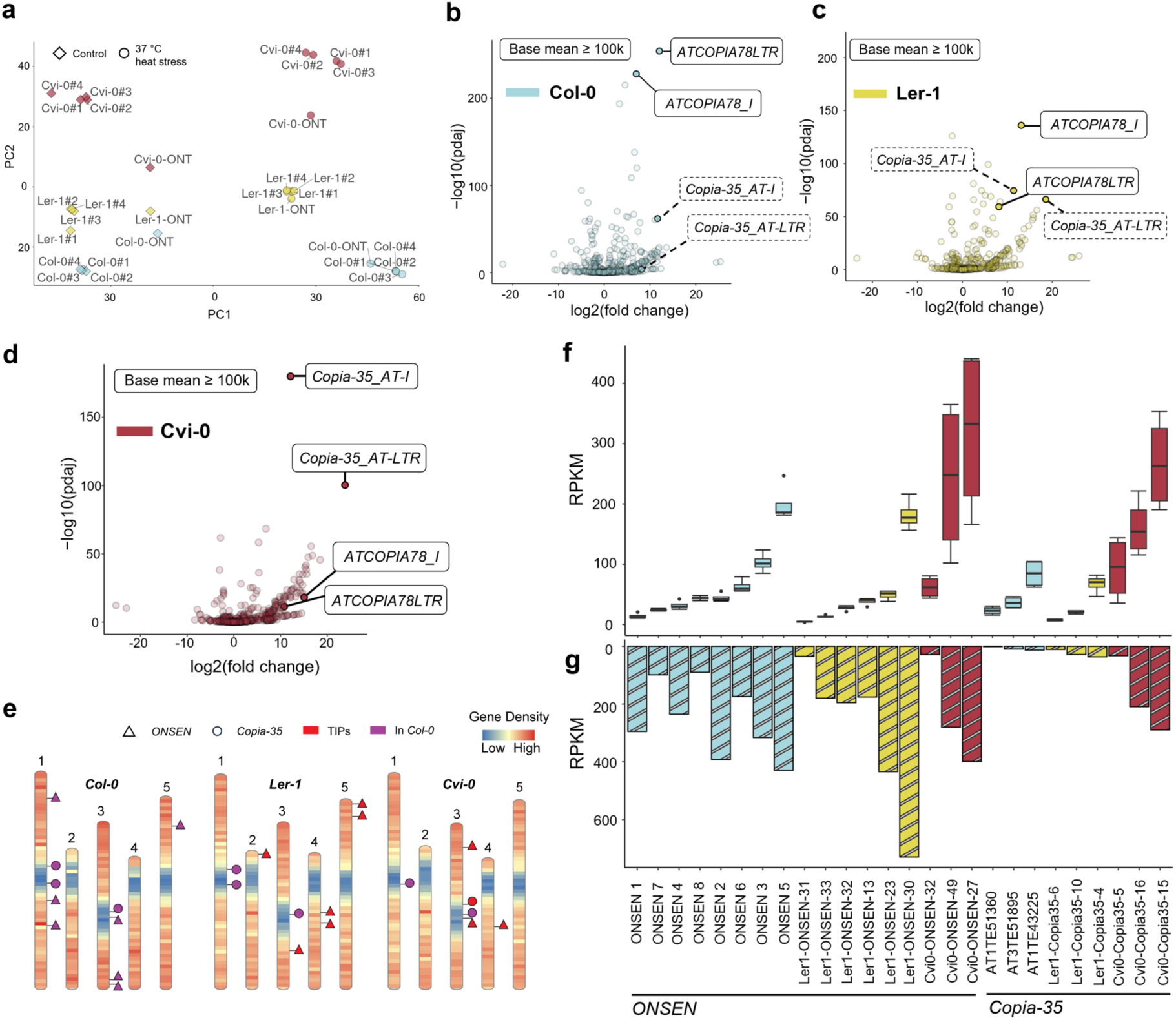
Expression of *ONSEN* and *Copia-35* a) PCoA analysis on gene expression in all sequenced samples. b-d) SalmonTE analysis with RNA-Seq data. Labeled consensus sequences in solid outlines represents candidates that have a base mean value that is greater than 100,000*. Copia-35* consensus sequences in b and c are labeled in dashed boxes due to below-cutoff base mean. e) Annotation of *ONSEN* and *Copia-35* full-length copies in the three accessions. Reference insertions and TIPs are marked. f) and g) Expression of TE per copy measured using RNA-Seq (four replicates) and ONT direct cDNA sequencing, respectively.

### Activity of heat-responsive TEs differs across accessions

We first aimed to identify TE candidates responsive to heat stress in each of the accessions. For this purpose, we used a consensus sequence-guided approach. Based on the library from Repbase (Bao et al., 2015), which contains 1,136 *A. thaliana* specific TE consensus sequences, we measured the transcriptional abundance of TEs in our RNA-Seq data using SalmonTE. Notably, in the Repbase library, the LTR and the internal consensus of LTR retrotransposons were constructed separately, enabling us to distinguish the expression of LTR vs. internal sequences. To reduce noise and to only focus on high-confidence TEs that would react to heat stress, we applied a stringent filter of log2 fold change ≥ 2 and padj ≤ 10^-10^, and a baseMean exceeding 100,000. We found that in all three accessions, the internal (*ATCOPIA78_I*) and the LTR (*ATCOPIA78LTR*) segments of *ONSEN* were significantly upregulated and with a high baseMean (Fig. 1b-d), confirming the robustness of *ONSEN*’s activation under heat stress. Importantly, in addition to the well-known case of *ONSEN*, we also found *Copia-35* in Cvi-0 that emerged as a top candidate, passing the same stringent filters as *ONSEN* (Fig. 1d). In Cvi-0, both *Copia-35_AT-I* and *Copia-35_AT-LTR* showed a high level of expression and even greater statistical significance when compared to the activation of the *ONSEN* family.

### Variations of expression of individual TE insertions

After assessing the global expression of *ONSEN* and *Copia-35* based on consensus sequences and RNA-Seq data, we combined the ONT direct cDNA-(ONT in short) and RNA-seq data to explore variations in expression among individual full-length TE copies of the same family. We first generated high confidence annotations of the two identified heat-responsive retrotransposon families *ONSEN* and *Copia-35* in all three accessions. In total, we identified six full-length *ONSEN* copies in Ler-1 and three in Cvi-0, as well as three full-length *Copia-35* copies in both accessions (Fig. 1e, Table S2 and 3). For Col-0, we adopted the TAIR10 annotation IDs for the full-length *ONSEN* and three *Copia-35* elements. However, we refined their annotations to include both LTRs. Interestingly, we found all full-length *ONSEN* insertions in Ler-1 and Cvi-0 to be polymorphic, representing TIPs (Fig. 1e). For *Copia-35*, one TIP was identified on chromosome 3 of Cvi-0, whereas all other full-length *Copia-35* insertions in Ler-1 and Cvi-0 were shared with Col-0 (Fig. 1e).

Subsequently, we aligned the RNA-Seq and ONT reads to their respective genomic assemblies, considering only uniquely mapped reads for downstream analysis. Overall, the pattern of expression levels was generally highly consistent between the RNA-seq and ONT for a given accession (e.g., *ONSEN 5* was the most expressed copy in Col-0, as was *ONSEN 30* in Ler-1, according to both datasets) (Fig. 1f-g). Both RNA-Seq and ONT revealed a significant variation of expression levels between individual *ONSEN* and *Copia-35* copies (Fig. 1f-g). In accordance with our consensus-based analysis, we found a specifically high activity of *Copia-35* in Cvi-0 compared to the other two accessions. Indeed, the least transcribed copy in Cvi-0, *Cvi0-Copia35-5*, reached expression levels resembling those of the most expressed *Copia-35* copies in the other two accessions. In addition, both RNA-seq and ONT datasets revealed similar expression levels of both TE-families in Cvi-0, with the highest expression level approximating 400 RPKM. Note that *ONSEN 7* was not included in further analyses as it harbors a large insertion, which together with its low expression level (Fig. 1e), suggests that this copy is not functional.

### ONT allows for a high-resolution profiling of *ONSEN* and *Copia-35*

Given the substantial differences in abundance of per-copy expression of *ONSEN* and *Copia-35*, we investigated the expression of individual copies in detail with ONT. Using the alignment of one of the most active and autonomous *ONSEN* copies (*ONSEN 1*) (Cavrak et al. 2014; Roquis et al. 2021), we found that, under heat stress, active full-length *ONSEN* copies have two transcription starting sites (TSS), namely S1 and S2, one within each of their LTRs (Fig 2a, Fig. S2-4). Moreover, we identified two transcription termination sites, E1 and E2. E1 is located just after the detected gag domain and E2 is situated at the 3’ LTR. A read from S1 to E2 thus represents a full-length mRNA that serves as a precursor for subsequent reverse transcription to *ONSEN*. Importantly, the RNA-Seq data failed to resolve the transcription starts and ends (Fig. 2b).

**Figure 2.**
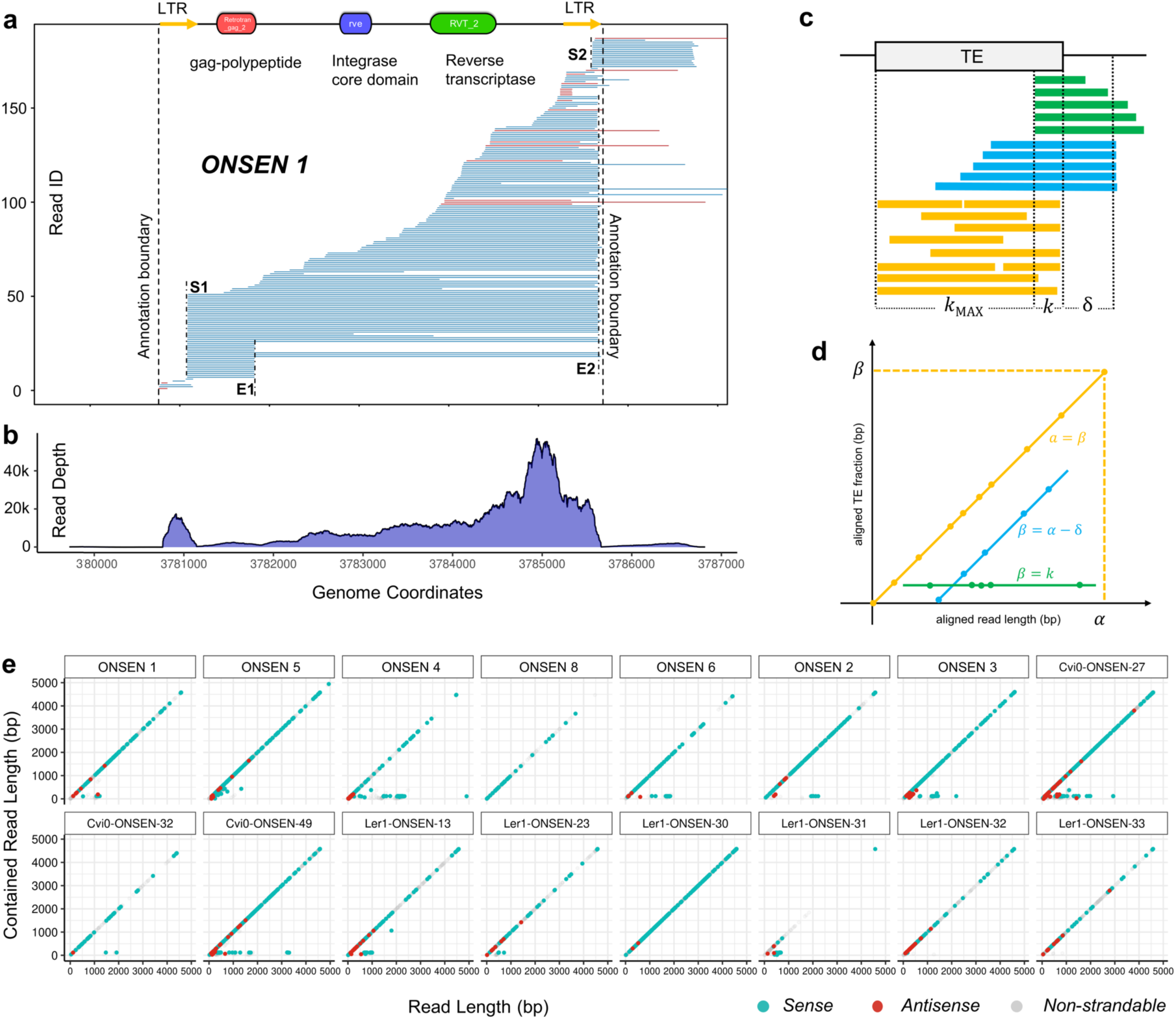
Transcriptional profile of *ONSEN* a) Long read alignment of *ONSEN 1*. Blue reads indicate matching orientation to the TE annotation (sense), while red reads indicate opposite orientation (antisense). b) Read depth of *ONSEN 1* in RNA-Seq data c-d) Principles of an aligned TE fraction vs read length plot. c) Read-through reads of a TE annotation can be divided into four groups. 1) alignments that cover the entire TE annotation (purple). 2) alignments that are contained in the TE annotation (yellow). 3) alignments that start outside of a TE annotation. 4) alignments that start within the TE annotation. Aligned read length and aligned TE fraction length is denoted as α and β respectively. The symbol δ indicates the distance between a transcription starting site outside of a TE annotation and the TE. The symbol k indicates the distance between the transcription starting site to one end of the TE. d) Example of a Transposon-Read Alignment Length Analysis (TRALA) plot, in which α is plotted against β e) TRALA plot of 16 full length *ONSEN* copies.

We also found that the 5’ LTR acts as a more dominant promoter than the 3’ LTR driving the selective expression of the gag-polypeptide or of the entire element, respectively. To quantify the difference in strength, we counted the number of reads from S1 and S2 for active *ONSEN* copies of all accessions (Table S2). We assumed that reads with starting sites between S1 and S2 were also transcribed from S1. This assumption was based on the rationale that many mRNA molecules were not fully sequenced to their 5’ ends, as suggested from the continuous distribution of reads across the entire elements (Fig. 2a), likely due to limitations of the reverse transcriptase during ONT library preparation. We found that the 5’ LTR accounts for 71.3% to 100% of the *ONSEN* transcripts, except for *ONSEN 4*, where the 3’ LTR accounts for 71.2% of the total transcription.

To assess the global variations of full-length *ONSEN* copies, we implemented a graphical analysis by plotting the aligned read length of an ONT read against the length covered by a TE annotation (Fig. 2c, d), which we refer to as Transposon-Read Alignment Length Analysis (TRALA) plot (Fig. 2d). As aforementioned, for most *ONSEN* copies, reads were initiated from S1 and therefore contained in the annotation, appearing as dots on the diagonal line. However, *ONSEN 4*, *Cvi0-ONSEN-27*, and *Cvi0-ONSEN-49* form a horizontal line at the bottom due to substantial amounts of reads initiated from S2, hence directly driving the expression of their flanking regions. Moreover, the TRALA plots revealed differences in the abundance of antisense transcription substantiating the expressional diversity among individual *ONSEN* copies (Fig. 2e).

We found that, like *ONSEN*, when exposed to heat stress, full-length copies of *Copia-35* show a continuous distribution of reads and have TSS S1 and S2 within each of their LTRs (Fig. 3a, Fig. S5-7). In contrast to *ONSEN*, we identified three termination sites: E1, E2, and E3 in *Copia-35*. E1 is located between the 5’ LTR and the gag-polypeptide, E2 is between the integrase and reverse transcriptase domain, while E3 lies at the 3’ LTR. Hence, a read from S1 to E3 represents a full-length mRNA that serves as a precursor for subsequent reverse transcription to *Copia-35* cDNA. As shown for the most active *Copia-35* copy (*Cvi0-Copia35-15*) and in contrast to the ONT data, RNA-Seq again failed to identify the transcription start and end points (Fig. 3b). Notably, the high-resolution provided by the ONT data also revealed that some of the reads aligning to *Copia-35* were spliced between S1 and E1 (Fig. 3a, Fig. S4-6).

**Figure 3.**
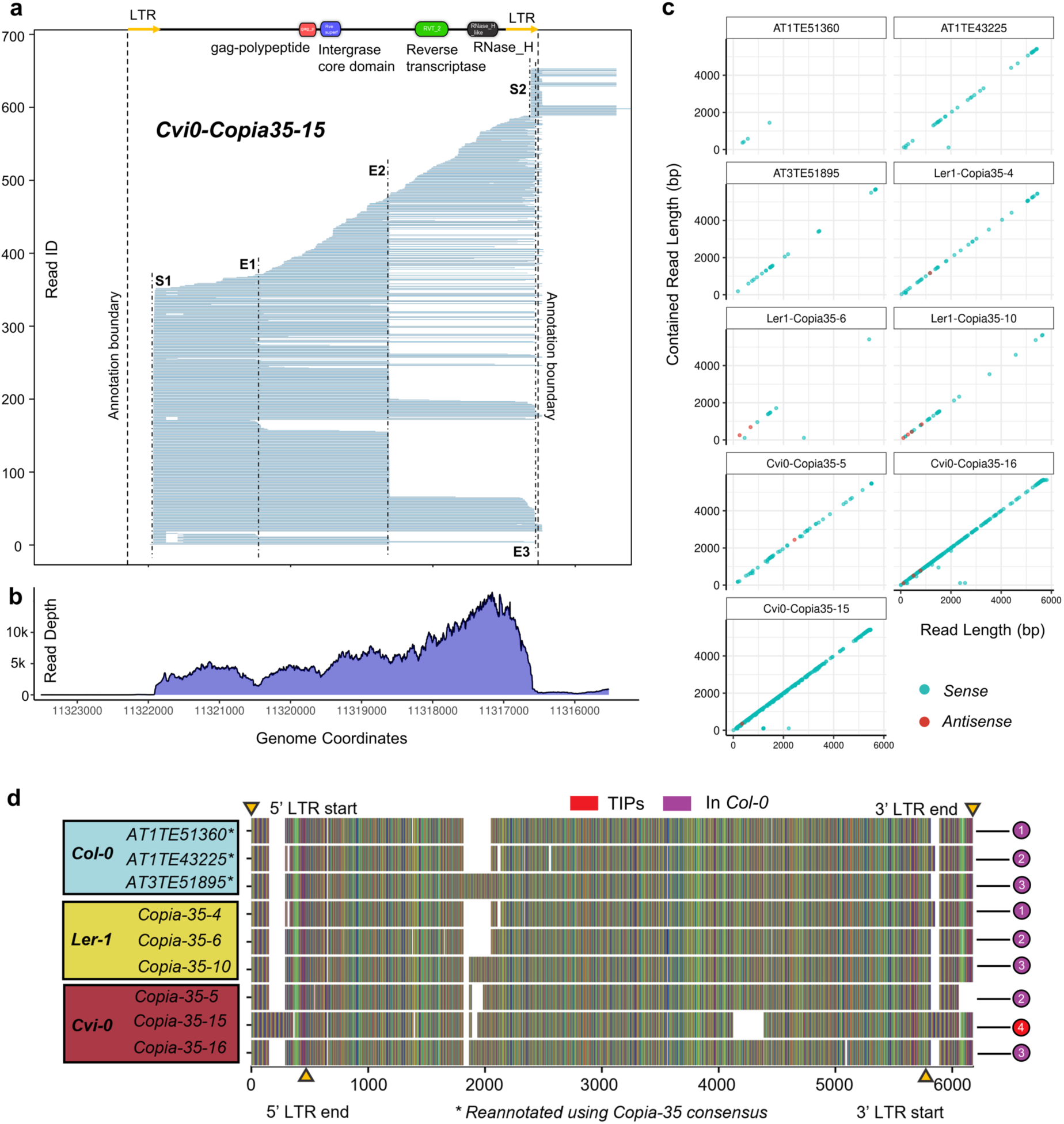
Expression of *Copia-35* profiled with long reads. a) Long reads alignment of *Cvi0-Copia35-15*. Blue reads indicate matching orientation to the TE annotation (sense). b) Read depth of *Cvi0-Copia35-15* in RNA-Seq data. c) TRALA plot of nine full length *Copia-35* copies d) Alignment of full length *Copia-35* copies in Col-0, Ler-1, and Cvi-0. LTR boundaries are marked by yellow triangles. Col-0 copies are numbered 1-3; numbers on sequences of other accessions correspond to these Col-0 copies conserved among accessions. The Cvi-0 TIP is labeled as copy No. 4.

The TRALA plot of all nine *Copia-35* copies revealed that most reads are contained within the *Copia-35* annotations (Fig. 3c), with the exception of *Cvi0-Copia35-15* and *Cvi0-Copia35-16*, which both show the existence of read-through transcripts. In addition to substantial differences between the number of transcripts per copy, the dots on the diagonal line in the TRALA plots of most *Copia-35* copies in Col-0 and Ler-1 contained large gaps, suggesting that not the entire length of the element is transcribed. To investigate whether obvious structural differences were responsible for this discrepancy between copies, we aligned all full-length *Copia-35* elements. We found that despite having greater expression, the full-length copies in Cvi-0 exhibited no major structural differences compared to copies in Ler-1 and Col-0 (Fig. 3d). For example, *Cvi0-Copia35-16* and *Ler1-Copia35-10* showed different expression levels under heat stress, but were identical in terms of structure, except for a small deletion in *Cvi0-Copia35-16* at around 5000 bp. Notably, we observed that the most active copy *Cvi0-Copia-35-15* that is also a TIP carried an insertion in both its LTRs.

### Both *ONSEN* and *Copia-35* confer heat responsiveness to their flanks

It is well established that full-length *ONSEN* elements can trigger the expression of adjacent genes under heat-stress (Ito et al. 2011; Roquis et al. 2021), a pattern we confirmed in our RNA-Seq data. Among the seven full-length *ONSEN* copies in Col-0, three caused the upregulation of both their 5’ and 3’ flanking genes, and a fourth triggered the upregulation of the 3’ genes only (Fig. 4a).

**Figure 4.**
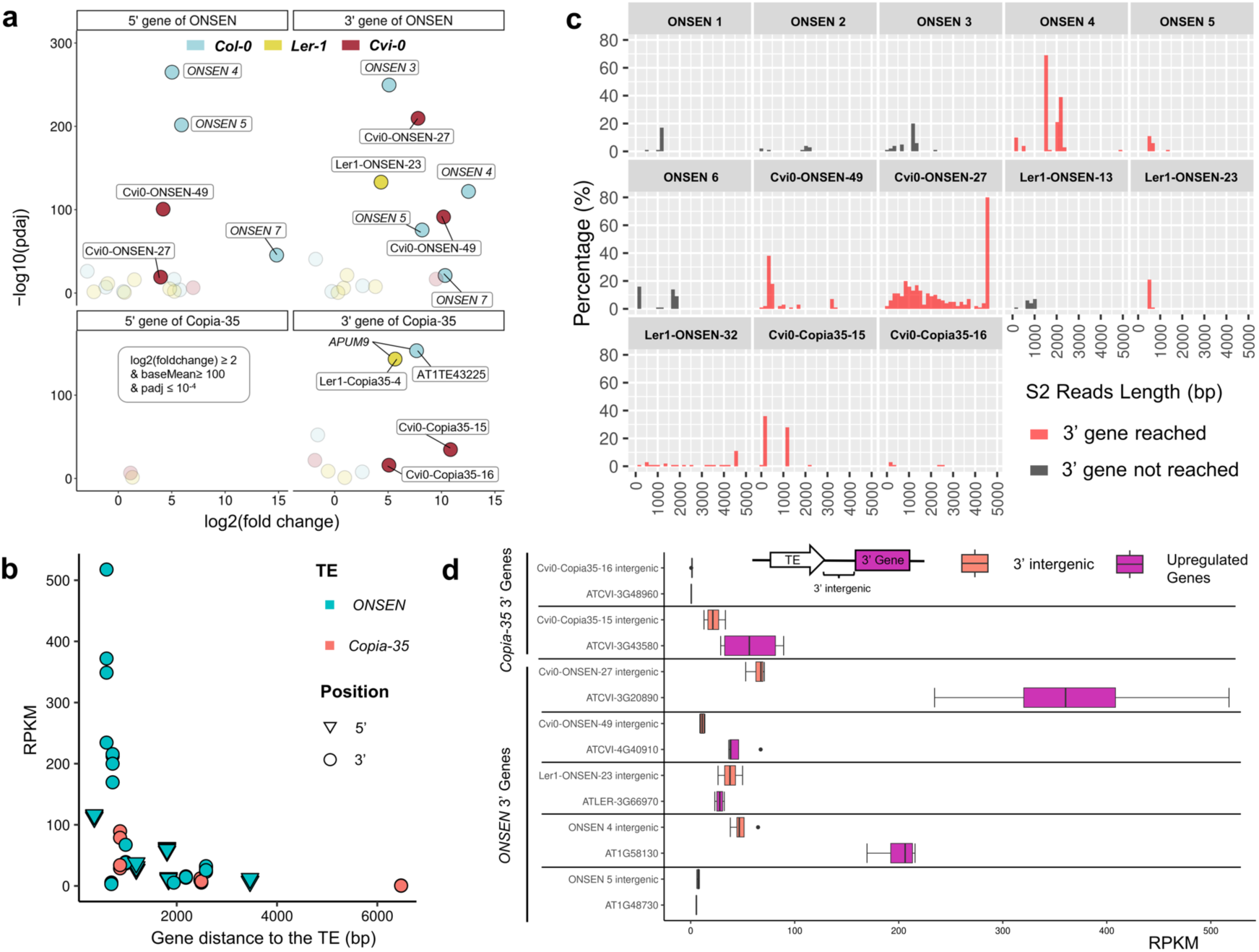
Upregulation of *ONSEN* and *Copia-35* flanking genes. a) Volcano plots highlighting genes adjacent to *ONSEN* and *Copia-35* with the following criteria: log2(fold change) ≥ 2, baseMean ≥ 100, and padj ≤ 10^-4^. Highlighted genes are labeled with the names of their corresponding TE. b) Relationship between a gene’s distance to its corresponding TE and RPKM. *ONSEN* genes are depicted in cyan, and *Copia-35* genes in coral. 5’ flanking genes (triangles) and 3’ flanking genes (circles) are denoted. c) Length distribution of S2 reads across 13 TEs. S2 reads reaching their 3’ genes are marked in coral. d) A comparison of the RPKM of the 3’ intergenic regions (located between the TE and its 3’ gene) against the RPKM of the corresponding upregulated 3’ gene, as described in a. Only upregulated 3’ genes reached by reads originating in the S2 of the TE are included in the analysis.

This pattern was also observed with two copies in Cvi-0 with the upregulation of flanking genes on both sides of *Cvi-0-ONSEN-27* and *Cvi-0-ONSEN-49* showing a log2fold change > 2 and padj < 10^-4^ (Fig. 4a), while in Ler-1, this was only observed for *Ler1-ONSEN-23* in the 3’ direction.

Since we found *Copia-35* expression in all three accessions, we next investigated whether this, by analogy with *ONSEN*, also induced expression of its flanking genes. Our data confirmed that the expression of *Cvi0-Copia35-15 and Cvi0-Copia35-16*, two predominantly expressed copies in Cvi-0, led to a significant upregulation of their 3’ flanking genes (Fig. 4a). Notably, while *Cvi0-Copia35-16* was shared between the three accessions (Fig. 3d) the upregulation of its 3’ gene was only observed in Cvi-0.

To determine whether the distance between the TE and the flanking genes could explain the observed patterns in Fig. 4a, we further plotted the distance between each gene and its associated TE against the gene’s RPKM. We uncovered a localized effect of TE-mediated gene activation under heat stress with closer genes showing a stronger heat response (Fig. 4b). To test whether the upregulation of flanking genes could also be explained by the detected read-out transcription from the 3’-LTR of some TE copies (Fig. 2a, Fig. 3a, Fig. S1-S6 and Table S2), we plotted the length of all S2 reads of TE copies that exhibit transcription from their 3’ LTR (Fig. 4c). For most copies, the length of S2 reads ranged between 0-2 kb. However, for some insertions we found that S2 reads were spanning up to 4.5 kb of the flanking region, even reaching the 3’ gene in seven cases (Fig 4d). To assess the importance of those reads in driving gene expression, we sought to quantify the relative transcription level of the intergenic region between the TE and the 3’ flanking gene (Table S2-3). This analysis showed that the expression of the intergenic region was either similar or lower than the actual gene expression. We further noted that the transcription of highly expressed flanking genes such as *AT1G58130* and *ATCVI-3G20890* was independent from the abundance of reads aligning to the flanking region (Fig. 4d), suggesting that the cis-regulatory effect of the TE is the main driver of their heat response.

Among the genes that were solely upregulated by the cis-regulatory effect of the TE (Fig. 4a, Table S2-3), we detected *APUM9,* a well-characterized gene that plays an important role in development (Xiang et al. 2014; Hristova et al. 2015). Indeed, *APUM9* was highly expressed in response to heat in Col-0 and Ler-1 but not in Cvi-0, where the *Copia-35* insertion was missing (Fig. 3d, Fig. 4a). Because the transcriptional changes of *APUM9* under heat stress may have phenotypic consequences and thus play a role in adaption, we further determined how frequently this TAP of *Copia-35* in the flanking region of *APUM9* occurred in natural accessions. After validating our approach using the available PacBio assemblies (Fig. S7, Table S4), we screened genomic reads of 1030 available accessions for the presence of this copy. Overall, we detected TAPs in 340 accessions, belonging to all genetic groups of *A. thaliana* (Fig. 5a). Surprisingly, TAPs were found in accessions geographically close to those carrying the *Copia-35* insertion at the *APUM9* locus.

**Figure 5.**
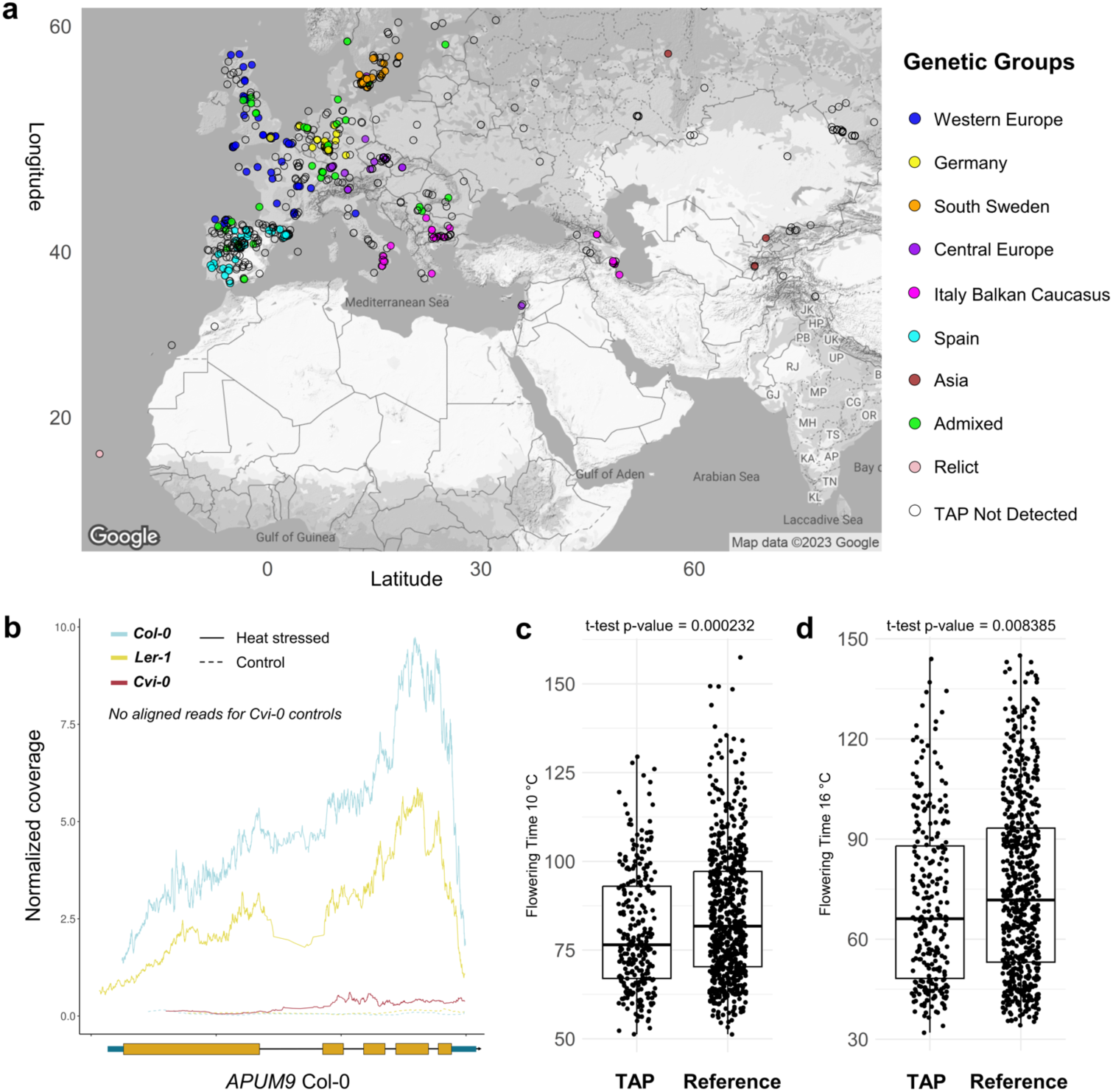
*APUM9* locus and flowering time of natural accessions of *A. thaliana*. a) Distribution map of the *Copia-35* TAP at the *APUM9* locus, with accessions color-coded by genetic group. b) Normalized RNA-Seq coverage for the *APUM9* gene across three accessions. Solid lines represent heat-stressed samples, while dashed lines represent controls. Normalized coverage is averaged over four replicates. Average flowering time at 10°C (**c**) and 16°C (**d**) depending on the detection of a *Copia-35* TAP at the *APUM9* locus. Reference indicates no TAP was detected.

Since our analysis showed that the expression of *APUM9* under heat stress was clearly associated with the presence of *Copia-35* (Fig. 4a, Fig. 5b) and knowing that *APUM9* is involved in regulating flowering time (Nyikó et al. 2019), we tested the possibility that the presence of *Copia-35* may affect this important trait when plants are exposed to different temperatures. By analyzing publicly available data, we found significant differences of flowering time at 10 (FT10, p-value < 0.001) and 16 °C (FT16, p-value < 0.01) depending on the presence of *Copia-35* in the flanking region of *APUM9* (Fig. 5c, d).

## Discussion

TE activity is an important source of transcriptional novelty (Rebollo et al. 2012) and a major driver of genome evolution. The genetic diversity arising from TE mobility has been documented in wild plants, including *A. thaliana* (Quadrana et al. 2016; Baduel et al. 2021) and *Brachypodium distachyon* (Stritt et al. 2020), as well as in crops like rice (Huang et al. 2008; Carpentier et al. 2019; Castanera et al. 2021), maize (Stitzer et al. 2021), and wheat (Wicker et al. 2022). While ONT long read sequencing has recently been shown to be effective to study TE expression in *Arabidopsis* mutants impaired for TE silencing (Panda and Slotkin 2020; Berthelier et al. 2023), the availability of high-quality assemblies now makes it possible to investigate the diversity of individual, highly similar TEs in multiple natural accessions of the same species. Using heat as an abiotic stress, our analysis revealed multiple layers of significant expressional diversity linked to stress-inducible TEs in *A. thaliana*.

Besides confirming the heat-responsiveness of the well-studied *ONSEN* family, the use of three different natural genetic backgrounds allowed for the in-depth characterization of *Copia-35*, a second retrotransposon family with an increased activity under heat stress. Despite sharing heat as environmental trigger, our data revealed striking differences between both families. Indeed, while none of the *ONSEN* copies is conserved between all three accessions, we only detected one TIP of *Copia-35* in the relict accession Cvi-0. These findings support the view that *ONSEN* is highly dynamic (Baduel et al. 2021), and could indicate a reduced mobility of *Copia-35* in Ler-1 and Col-0 compared to Cvi-0. This argument is further strengthened by the fact that *Copia-35* elements in Col-0 are lacking the ability to transpose, pointing towards a non-autonomous nature in this accession (Pietzenuk et al., 2016).

In response to heat treatment, both ONT and RNA-seq data showed that the transcription of *Copia-35* was relatively low in Col-0 and Ler-1 but reached high expression levels, similar to those of *ONSEN*, in Cvi-0. Our ONT data further confirmed the presence of full-length transcripts that could serve as a template for the reverse transcription resulting in the transposition of *Copia-35* in Cvi-0. These results show that the genome of Cvi-0 harbors two independent and potentially mobile TE families, synchronically activated by the same environmental trigger. Whether additional factors, such as specific insertion preferences as observed for *ONSEN* (Quadrana et al. 2019; Roquis et al. 2021) or their epigenetic regulation by different pathways, are defining separate ‘niches’ (Kidwell and Lisch, 1997; Venner et al., 2009) allowing for a coexistence of both families, remains to be elucidated.

The strong variation in the activity of *Copia-35,* which is equally abundant in all three accessions but differentially expressed, is in line with previous work (Marí-Ordóñez et al. 2013; Thieme et al. 2017; Nozawa et al. 2022), and suggests that factors other than copy number determine the overall activity of a TE-family. For instance, *Copia-35* expression increases in mutants deficient in epigenetic silencing (Yokthongwattana et al. 2010) while the loss of RdDM alone (i.e without abiotic stress), does not activate *ONSEN* (Ito et al. 2011), highlighting differences in the factors governing the activities of both families. Notably, recent work showed that natural variations in the strength of epigenetic silencing under heat stress leads to increased activation of *ONSEN* in the Kyoto accession that displays reduced methylation in the CHH context (Nozawa et al. 2022). In this regard, it is noteworthy that the relic accession Cvi-0 that displayed a high activity of both TEs in our study is globally hypomethylated compared to Col-0 (Kawakatsu et al. 2016).

The high resolution of the ONT data also revealed striking qualitative expressional differences between both families. Most importantly, we revealed the presence of an additional transcription termination site for *Copia-35* compared to *ONSEN.* This could imply mechanistic variations in the lifecycle of the two families. Analogous to retroviruses, LTR-RT require specific amounts of the structural GAG nucleocapsid, the catalytic polyprotein and the full-length transcript that serves as a template for reverse transcription to complete their lifecycle (Schulman 2013). Besides mechanisms affecting translation (Clare et al. 1988; Matthews et al. 1997; Havecker and Voytas 2003) subgenomic TE expression and splicing resulting in different transcript pools underly the fine-tuning of retrotransposon protein abundances (Chang et al. 2013). The role of alternative splicing is perfectly illustrated by its importance for regulating protein abundances of the *Arabidopsis Copia*-type retrotransposon *EVADÉ* (Oberlin et al. 2017). Our work, however, paints a more nuanced picture. While we detected the presence of a few spliced transcripts produced by *Copia-35*, our ONT analysis suggests the presence of short subgenomic transcripts that may indicate that the diverse RNA pools needed to complete the TE-lifecycle are obtained using a splicing-independent mechanism. These findings therefore open new avenues for elucidating the fundamental processes of plant retrotransposon mobility. This is particular crucial, because while *ONSEN* has been studied in detail (Ito et al. 2011; Cavrak et al. 2014; Thieme et al. 2017; Baduel et al. 2021) our current mechanistic understanding of plant TEs is overwhelmingly based on studies using few genetic backgrounds, and in the case of heat-responsive TEs, mainly on Col-0.

The influence of TEs on the expression of their flanking regions is well-documented (Butelli et al. 2012; Makarevitch et al. 2015; Rech et al. 2022). Here, we confirmed that *ONSEN* mediates a heat-dependent upregulation of flanking regions (Ito et al. 2011; Roquis et al. 2021) and further revealed that *Copia-35* can also confer heat-responsiveness to its neighboring genes, in addition to the previously reported *APUM9* locus in Col-0 (Pietzenuk et al. 2016), in multiple accessions. The ONT data further allowed us to unambiguously discriminate between read-out transcription and the indirect upregulation of genes via the *cis*-regulatory effect mediated by the recruitment of the transcription machinery to the TE (Zhao et al. 2018; Fagny et al. 2020; Deneweth et al. 2022). The formation of TE-gene fusion transcripts is a common phenomenon in *Arabidopsis* (Lockton and Gaut 2009; Berthelier et al. 2023) and we indeed detected read-out transcription originating from the 3’ LTRs of both *ONSEN* and *Copia-35* TE families under heat-stress. However, our data suggests that the *cis*-regulatory effect is the main driver of TE-mediated expression of the flanking genes. Interestingly, one of the genes that has previously been shown to be affected by *Copia-35* (Pietzenuk et al. 2016) is *APUM9*, which is involved in early embryonic development, with a putative role in basal heat tolerance (Nyikó et al. 2019). In addition, an overexpression of *APUM9* results in abnormal leaf morphology and a delayed flowering phenotype (Nyikó et al. 2019). Despite its importance in development, the natural diversity of the *APUM9* locus and more specifically the role of *Copia-35* in driving its expression under heat stress, had not been studied across multiple natural accessions. Our data revealed that on a population scale, accessions without the *Copia-35* insertion at the *APUM9* locus tend to flower earlier. The timing of flowering is crucial for a population to survive. Despite their selfish nature, major (epi)genetic effects linked to transposition events are generally viewed as a driving force of plant evolution (Lisch 2013), capable of facilitating rapid adaptation (Hof et al. 2016; Thieme et al. 2022), and the link between transposition and modulation of flowering time in *A. thaliana* has been suggested previously (Thieme et al. 2017; Quadrana et al. 2019; Baduel et al. 2021). Flowering time is a complex trait driven by multiple loci with small quantitative effects (Kinoshita and Richter 2020). The fact that heat triggers the upregulation of *Copia-35*, resulting in an activation of *APUM9*, and that the experimentally induced overexpression of *APUM9* in Col-0 results in delayed flowering (Nyikó et al. 2019), indeed indicates a quantitative effect of this insertion on flowering time.

Overall, our study revealed a great expressional diversity linked to heat-responsive LTR-retrotransposons in *A. thaliana*. These findings strongly advocate for the use of ONT in studies aiming at understanding both the fundamental mechanisms of LTR-retrotransposon mobility and their adaptive consequences across multiple natural accessions. With the increasing availability of high-quality genomes, similar studies should soon allow us to drastically improve our understanding of the role of TEs in plants that are densely packed with TEs.

## Materials and Methods

### Heat stress experiments, RNA extractions and sequencing

Seeds of Col-0, Ler-1 and Cvi-0 were first stratified on ½ Murashige and Skoog (MS) plates for 7 days at 4°C and then grown under controlled conditions (16 h light at 24°C, 8 hours dark at 22°C) in a Aralab 600 growth chamber (Rio de Mouro, Portugal). After 7 days of growth, plants were heat-stressed at 37°C for 24 h and 16 h light in a second Aralab 600 growth chamber. Seedlings from control and heat treatment were sampled simultaneously at the end of the stress period. For the ONT direct cDNA sequencing, 20 seedings per accession per treatment were pooled together for mRNA extraction using oligo-dT beads (#61011) (Thermo Fisher Scientific, Waltham, USA). The Functional Genomic Centre at Zürich performed library preparation and sequencing. Final cDNA libraries were sequenced on ONT Flow Cells (R 9.4.1) (Oxford, UK).

For the illumina RNA-Seq samples, plants were grown and stressed under the same conditions. Four biological replicates (pools of at least nine seedlings) per condition for each accession were extracted using the QIAGEN RNeasy plant mini kit (#74904) (Venlo, Netherlands). Novogene UK performed the library prep and sequencing.

### TE annotation

For *ONSEN*, full-length copies (Cavrak et al. 2014) were used to generate annotations using RepeatMasker (version 4.1.1) (repeatmasker.org) with the following options: -a -xsmall -gccalc - nolow. We only conducted the rest of the analysis on the remaining seven functional copies. In addition, TE consensus sequences of *A. thaliana* from RepBase28.03 (Bao et al. 2015) were used to annotate all other TEs using the same command. *ROMANIAT5* consensus sequence was reconstructed by Repbase in 2018 and its name was reverted to *Copia-35* (girinst.org/2018/vol18/issue9/Copia-35_AT-I.html). For clarity, this article abandoned the legacy name of *ROMANIAT5* and refers to the family as *Copia-35.* In the case of full-length copies of *Copia-35* in Col-0, we adopted their TAIR10 names, *AT1E51360*, *AT1E43225* and *AT3TE51895*, even after reannotation. For the remaining accessions, the elements were named based on the format: Accession-TE family-Annotation ID. NCBI conserved domain search (CDD v3.20) (Lu et al. 2019) was used to annotate protein domains in TE sequences.

### RNA-seq analysis

Fastp (version 0.23.2) (Chen 2023) was used to trim adapters and remove low complexity reads using the following options: --qualified_quality_phred 15 --unqualified_percent_limit 40 -- n_base_limit 10 --low_complexity_filter --correction --detect_adapter_for_pe -- overrepresentation_analysis --dedup --dup_calc_accuracy 6. Ribosomal RNA was then removed using bbduk.sh (version 39.01) from the BBTools suite (<u>sourceforge.net/projects/bbmap/</u>) with the options k=31 hdist=1.

Cleaned reads were then mapped to their respective genome assemblies using STAR (version 2.7.10b) (Dobin et al. 2012) with options: --alignIntronMax 5000 –outFilterMultimapNmax 100 –winAnchorMultimapNmax 100. The genome assembly and gene annotation of Col-0 (release 10) was downloaded from the Arabidopsis Information Resource (TAIR) (Berardini et al. 2015). The genome assemblies and gene annotation of Ler-1 and Cvi-0 were downloaded from the 1001 genomes webpage (Jiao and Schneeberger 2020).

We employed RPKM (Reads Per Kilobase of transcript, per Million mapped reads), a commonly used unit of measurement to quantify gene and TE expression levels and normalize the expression levels across replicates. Pair-ended fragments were counted using featureCounts (Liao et al. 2013) against the TE or gene annotations, with the following options: -B -p -P -O.

Cleaned RNA-Seq data were also analyzed by SalmonTE (version 0.4) (Jeong et al. 2017) to measure global expression of TEs. The *A. thaliana* TE consensus library was downloaded from Repbase (version 28.03.2023) (Bao et al., 2015) and used as the custom library for SalmonTE. Default options of SalmonTE’s “quant” and “test” program were used to quantify expression and perform statistical analyses.

### Basecalling and mapping of ONT data

Basecalling was performed on the passed fast5 files using Guppy (version 6.1.2) with default options. Guppy is developed by ONT and available via their community website (<u>community.nanoporetech.com</u>). Stranding was then directly performed on the passed output from basecalling using Pychopper (version 2.5.0) (github.com/epi2me-labs/pychopper). Primer configuration for stranding was set to "+:SSP,-VNP|-:VNP,-SSP’’ and rescued reads were not used. Porechop (version 0.2.4) (<u>github.com/rrwick/Porechop</u>) was then used to remove sequencing adapters from ONT reads. Finally, ONT reads were mapped to their respective genome assemblies using minimap2 (version 2.24) (Li 2018) with options -ax splice -uf -k14.

### Mapping of whole-genome sequencing (WGS) data

The WGS data of 1,135 *A. thaliana* accessions was downloaded from the National Center for Biotechnology Information Sequence Read Archive (NCBI SRA) under project PRJNA273563 (Alonso-Blanco et al. 2016). Fastp (version 0.23.2) (Chen 2023) was used to trim adapters and remove low complexity reads using the following options: --qualified_quality_phred 15 -- unqualified_percent_limit 40 --n_base_limit 10 --low_complexity_filter --correction -- detect_adapter_for_pe --overrepresentation_analysis --dedup --dup_calc_accuracy 6. BWA-MEM (version 0.7.17) (Li 2013) was used to map the genomic reads to the *APUM9* locus of Col-0.

### TAP detection at the *AMUP9* locus

Data retrieved from the 1001genomes project (Alonso-Blanco et al. 2016) was used to screen for TE Absence Polymorphisms (TAPs) at the *APUM9* locus. BWA-mem (version 0.7.17) (Li 2013) and detettore (version 2.0.3) (github.com/cstritt/detettore) was used in tandem to first map the reads, and then perform TAP calling using default options.

### Flowering time analysis

Flowering time at 16°C (FT16) and 10°C (FT10) recorded by the 1001genomes project (Alonso-Blanco et al. 2016) were used to test the association between the number of TAPs and flowering time.

## Data access

Raw RNA-seq and base-called ONT data were uploaded to the European Nucleotide Archive (ENA) under project PRJEB64476. The scripts used for the statistics and figure generation were deposited into https://github.com/GroundB/Natural-diversity-of-heat-induced-transcription-of-retrotransposons-in-Arabidopsis-thaliana.

## Acknowledgments

The author would like to thank the Functional Genomic Center Zurich for processing the samples and Dr Emmanuelle Botté (https://manuscribe.com.au) for professional editing of the manuscript.

This work was supported by the University of Zurich Research Priority Programs (URPP) Evolution in Action (M.T. and A.C.R) and the Schweizerischer Nationalfonds zur Förderung der Wissenschaftlichen Forschung. Grant Number: 31003A_182785 (A.C.R and W.X).

## Author contributions

M.T. and A.C.R. conceived the study; W.X. and M.T. conducted experiments; W.X. analyzed the data; W.X. and M.T. wrote the paper with contributions from A.C.R. A.C.R. secured funding. All authors approve the paper.

## References

Alonso-Blanco C, Andrade J, Becker C, Bemm F, Bergelson J, Borgwardt KM, Cao J, Chae E, Dezwaan TM, Ding W et al. 2016. 1,135 Genomes Reveal the Global Pattern of Polymorphism in *Arabidopsis thaliana*. Cell 166: 481–491.

Baduel P, Leduque B, Ignace A, Gy I, Gil J, Loudet O, Colot V, Quadrana L. 2021. Genetic and environmental modulation of transposition shapes the evolutionary potential of Arabidopsis thaliana. Genome Biology 22: 138.

Bao W, Kojima KK, Kohany O. 2015. Repbase Update, a database of repetitive elements in eukaryotic genomes. Mobile DNA 6: 11.

Berardini TZ, Reiser L, Li D, Mezheritsky Y, Muller R, Strait E, Huala E. 2015. The arabidopsis information resource: Making and mining the “gold standard” annotated reference plant genome. genesis 53: 474–485.

Berthelier J, Furci L, Asai S, Sadykova M, Shimazaki T, Shirasu K, Saze H. 2023. Long-read direct RNA sequencing reveals epigenetic regulation of chimeric gene-transposon transcripts in Arabidopsis thaliana. Nature Communications 14: 3248.

Butelli E, Licciardello C, Zhang Y, Liu J, Mackay S, Bailey P, Reforgiato-Recupero G, Martin C. 2012. Retrotransposons Control Fruit-Specific, Cold-Dependent Accumulation of Anthocyanins in Blood Oranges The Plant Cell 24: 1242–1255.

Carpentier MC, Manfroi E, Wei FJ, Wu HP, Lasserre E, Llauro C, Debladis E, Akakpo R, Hsing YI, Panaud O. 2019. Retrotranspositional landscape of Asian rice revealed by 3000 genomes. Nat Commun 10: 24.

Castanera R, Vendrell-Mir P, Bardil A, Carpentier M-C, Panaud O, Casacuberta JM. 2021. Amplification dynamics of miniature inverted-repeat transposable elements and their impact on rice trait variability. The Plant Journal 107: 118–135.

Cavrak VV, Lettner N, Jamge S, Kosarewicz A, Bayer LM, Mittelsten Scheid O. 2014. How a Retrotransposon Exploits the Plant’s Heat Stress Response for Its Activation. PLoS Genet 10: e1004115.

Chang W, Jääskeläinen M, Li SP, Schulman AH. 2013. BARE retrotransposons are translated and replicated via distinct RNA pools. PLoS One 8: e72270.

Chen S. 2023.Ultrafast one-pass FASTQ data preprocessing, quality control, and deduplication using fastp. *iMeta* 2: e107.

Clare JJ, Belcourt M, Farabaugh PJ. 1988. Efficient translational frameshifting occurs within a conserved sequence of the overlap between the two genes of a yeast Ty1 transposon. Proceedings of the National Academy of Sciences 85: 6816–6820.

Deneweth J, Van de Peer Y, Vermeirssen V. 2022. Nearby transposable elements impact plant stress gene regulatory networks: a meta-analysis in A. thaliana and S. lycopersicum. BMC Genomics 23: 18.

Dobin A, Davis CA, Schlesinger F, Drenkow J, Zaleski C, Jha S, Batut P, Chaisson M, Gingeras TR. 2012. STAR: ultrafast universal RNA-seq aligner. Bioinformatics 29: 15–21.

Fagny M, Kuijjer ML, Stam M, Joets J, Turc O, Rozière J, Pateyron S, Venon A, Vitte C. 2020. Identification of Key Tissue-Specific, Biological Processes by Integrating Enhancer Information in Maize Gene Regulatory Networks. Front Genet 11: 606285.

Havecker ER, Voytas DF. 2003. The soybean retroelement SIRE1 uses stop codon suppression to express its envelope-like protein. EMBO Rep 4: 274–277.

Hof AEvt, Campagne P, Rigden DJ, Yung CJ, Lingley J, Quail MA, Hall N, Darby AC, Saccheri IJ. 2016. The industrial melanism mutation in British peppered moths is a transposable element. Nature 534: 102–105.

Hristova E, Fal K, Klemme L, Windels D, Bucher E. 2015. HISTONE DEACETYLASE6 Controls Gene Expression Patterning and DNA Methylation-Independent Euchromatic Silencing Plant Physiology 168: 1298–1308.

Huang X, Lu G, Zhao Q, Liu X, Han B. 2008. Genome-wide analysis of transposon insertion polymorphisms reveals intraspecific variation in cultivated rice. Plant Physiol 148: 25–40.

Ito H, Gaubert H, Bucher E, Mirouze M, Vaillant I, Paszkowski J. 2011. An siRNA pathway prevents transgenerational retrotransposition in plants subjected to stress. Nature 472: 115–119.

Ito H, Yoshida T, Tsukahara S, Kawabe A. 2013. Evolution of the ONSEN retrotransposon family activated upon heat stress in Brassicaceae. Gene 518: 256–261.

Jeong H-H, Yalamanchili HK, Guo C, Shulman JM, Liu Z. 2017. An ultra-fast and scalable quantification pipeline for transposable elements from next generation sequencing data. In Biocomputing 2018, doi:doi:10.1142/9789813235533_0016 10.1142/9789813235533_0016, pp. 168–179. WORLD SCIENTIFIC.

Jeong HH, Yalamanchili HK, Guo C, Shulman JM, Liu Z. 2018. An ultra-fast and scalable quantification pipeline for transposable elements from next generation sequencing data. Pac Symp Biocomput 23: 168–179.

Jiao W-B, Schneeberger K. 2020. Chromosome-level assemblies of multiple Arabidopsis genomes reveal hotspots of rearrangements with altered evolutionary dynamics. Nature Communications 11: 989.

Jiao Y, Peluso P, Shi J, Liang T, Stitzer MC, Wang B, Campbell MS, Stein JC, Wei X, Chin C-S et al. 2017. Improved maize reference genome with single-molecule technologies. Nature 546: 524–527.

Jin Y, Tam OH, Paniagua E, Hammell M. 2015. TEtranscripts: a package for including transposable elements in differential expression analysis of RNA-seq datasets. Bioinformatics 31: 3593–3599.

Kawakatsu T, Huang S-sC, Jupe F, Sasaki E, Schmitz RJ, Urich MA, Castanon R, Nery JR, Barragan C, He Y, et al. 2016. Epigenomic Diversity in a Global Collection of *Arabidopsis thaliana* Accessions. Cell 166: 492–505.

Kinoshita A, Richter R. 2020. Genetic and molecular basis of floral induction in Arabidopsis thaliana. Journal of experimental botany 71: 2490–2504.

Lanciano S, Cristofari G. 2020. Measuring and interpreting transposable element expression. Nature Reviews Genetics 21: 721–736.

Li H. 2013. Aligning sequence reads, clone sequences and assembly contigs with BWA-MEM. arXiv preprint arXiv:13033997.

Li H. 2018. Minimap2: pairwise alignment for nucleotide sequences. Bioinformatics 34: 3094–3100.

Liao Y, Smyth GK, Shi W. 2013. featureCounts: an efficient general purpose program for assigning sequence reads to genomic features. Bioinformatics 30: 923–930.

Lisch D. 2013. How important are transposons for plant evolution? Nat Rev Genet 14: 49–61.

Lockton S, Gaut BS. 2009. The contribution of transposable elements to expressed coding sequence in Arabidopsis thaliana. J Mol Evol 68: 80–89.

Lu S, Wang J, Chitsaz F, Derbyshire MK, Geer RC, Gonzales NR, Gwadz M, Hurwitz DI, Marchler GH, Song JS et al. 2019. CDD/SPARCLE: the conserved domain database in 2020. Nucleic Acids Research 48: D265–D268.

Makarevitch I, Waters AJ, West PT, Stitzer M, Hirsch CN, Ross-Ibarra J, Springer NM. 2015. Transposable Elements Contribute to Activation of Maize Genes in Response to Abiotic Stress. PLoS Genet 11: e1004915.

Marí-Ordóñez A, Marchais A, Etcheverry M, Martin A, Colot V, Voinnet O. 2013. Reconstructing de novo silencing of an active plant retrotransposon. Nature Genetics 45: 1029–1039.

Masuda S, Nozawa K, Matsunaga W, Masuta Y, Kawabe A, Kato A, Ito H. 2016. Characterization of a heat-activated retrotransposon in natural accessions of *Arabidopsis thaliana*. Genes & Genetic Systems 91: 293–299.

Matthews GD, Goodwin TJ, Butler MI, Berryman TA, Poulter RT. 1997. pCal, a highly unusual Ty1/copia retrotransposon from the pathogenic yeast Candida albicans. J Bacteriol 179: 7118–7128.

Matzke MA, Mosher RA. 2014. RNA-directed DNA methylation: an epigenetic pathway of increasing complexity. Nature Reviews Genetics 15: 394–408.

Negi P, Rai AN, Suprasanna P. 2016. Moving through the Stressed Genome: Emerging Regulatory Roles for Transposons in Plant Stress Response. Front Plant Sci 7: 1448.

Nozawa K, Masuda S, Saze H, Ikeda Y, Suzuki T, Takagi H, Tanaka K, Ohama N, Niu X, Kato A et al. 2022. Epigenetic regulation of ecotype-specific expression of the heat-activated transposon ONSEN. Front Plant Sci 13: 899105.

Nyikó T, Auber A, Bucher E. 2019. Functional and molecular characterization of the conserved Arabidopsis PUMILIO protein, APUM9. Plant Molecular Biology 100: 199–214.

Oberlin S, Sarazin A, Chevalier C, Voinnet O, Marí-Ordóñez A. 2017. A genome-wide transcriptome and translatome analysis of Arabidopsis transposons identifies a unique and conserved genome expression strategy for Ty1/Copia retroelements. Genome Res 27: 1549–1562.

Panda K, Slotkin RK. 2020. Long-Read cDNA Sequencing Enables a “Gene-Like” Transcript Annotation of Transposable Elements. The Plant Cell 32: 2687–2698.

Pecinka A, Dinh HQ, Baubec T, Rosa M, Lettner N, Scheid OM. 2010. Epigenetic Regulation of Repetitive Elements Is Attenuated by Prolonged Heat Stress in Arabidopsis The Plant Cell 22: 3118–3129.

Pietzenuk B, Markus C, Gaubert H, Bagwan N, Merotto A, Bucher E, Pecinka A. 2016. Recurrent evolution of heat-responsiveness in Brassicaceae COPIA elements. Genome Biology 17: 209.

Quadrana L, Bortolini Silveira A, Mayhew GF, LeBlanc C, Martienssen RA, Jeddeloh JA, Colot V. 2016. The Arabidopsis thaliana mobilome and its impact at the species level. eLife 5: e15716.

Quadrana L, Etcheverry M, Gilly A, Caillieux E, Madoui M-A, Guy J, Bortolini Silveira A, Engelen S, Baillet V, Wincker P et al. 2019. Transposition favors the generation of large effect mutations that may facilitate rapid adaption. Nature Communications 10: 3421.

Rebollo R, Romanish MT, Mager DL. 2012. Transposable Elements: An Abundant and Natural Source of Regulatory Sequences for Host Genes. Annual Review of Genetics 46: 21–42.

Rech GE, Radío S, Guirao-Rico S, Aguilera L, Horvath V, Green L, Lindstadt H, Jamilloux V, Quesneville H, González J. 2022. Population-scale long-read sequencing uncovers transposable elements associated with gene expression variation and adaptive signatures in Drosophila. Nature Communications 13: 1948.

Roquis D, Robertson M, Yu L, Thieme M, Julkowska M, Bucher E. 2021. Genomic impact of stress-induced transposable element mobility in Arabidopsis. Nucleic Acids Research 49: 10431–10447.

Sanchez DH, Paszkowski J. 2014. Heat-Induced Release of Epigenetic Silencing Reveals the Concealed Role of an Imprinted Plant Gene. PLoS Genet 10: e1004806.

Schulman AH. 2013. Retrotransposon replication in plants. Current Opinion in Virology 3: 604–614.

Stitzer MC, Anderson SN, Springer NM, Ross-Ibarra J. 2021. The genomic ecosystem of transposable elements in maize. PLoS Genet 17: e1009768.

Stritt C, Wyler M, Gimmi EL, Pippel M, Roulin AC. 2020. Diversity, dynamics and effects of long terminal repeat retrotransposons in the model grass Brachypodium distachyon. New Phytologist 227: 1736–1748.

Sun L, Jing Y, Liu X, Li Q, Xue Z, Cheng Z, Wang D, He H, Qian W. 2020. Heat stress-induced transposon activation correlates with 3D chromatin organization rearrangement in Arabidopsis. Nature Communications 11: 1886.

Thieme M, Brêchet A, Bourgeois Y, Keller B, Bucher E, Roulin AC. 2022. Experimentally heat-induced transposition increases drought tolerance in Arabidopsis thaliana. New Phytologist **n/a**.

Thieme M, Lanciano S, Balzergue S, Daccord N, Mirouze M, Bucher E. 2017. Inhibition of RNA polymerase II allows controlled mobilisation of retrotransposons for plant breeding. Genome Biology 18: 134.

Tittel-Elmer M, Bucher E, Broger L, Mathieu O, Paszkowski J, Vaillant I. 2010. Stress-Induced Activation of Heterochromatic Transcription. PLoS Genet 6: e1001175.

Wicker T, Gundlach H, Spannagl M, Uauy C, Borrill P, Ramírez-González RH, De Oliveira R, Mayer KFX, Paux E, Choulet F et al. 2018. Impact of transposable elements on genome structure and evolution in bread wheat. Genome Biology 19: 103.

Wicker T, Sabot F, Hua-Van A, Bennetzen JL, Capy P, Chalhoub B, Flavell A, Leroy P, Morgante M, Panaud O et al. 2007. A unified classification system for eukaryotic transposable elements. Nature Reviews Genetics 8: 973–982.

Wicker T, Schulman AH, Tanskanen J, Spannagl M, Twardziok S, Mascher M, Springer NM, Li Q, Waugh R, Li C et al. 2017. The repetitive landscape of the 5100 Mbp barley genome. Mobile DNA 8: 22.

Wicker T, Stritt C, Sotiropoulos AG, Poretti M, Pozniak C, Walkowiak S, Gundlach H, Stein N. 2022. Transposable Element Populations Shed Light on the Evolutionary History of Wheat and the Complex Co-Evolution of Autonomous and Non-Autonomous Retrotransposons. Advanced Genetics 3: 2100022.

Wu C. 1995. Heat Shock Transcription Factors: Structure and Regulation. Annual Review of Cell and Developmental Biology 11: 441–469.

Xiang Y, Nakabayashi K, Ding J, He F, Bentsink L, Soppe WJ. 2014. Reduced Dormancy5 encodes a protein phosphatase 2C that is required for seed dormancy in Arabidopsis. Plant Cell 26: 4362–4375.

Yokthongwattana C, Bucher E, Caikovski M, Vaillant I, Nicolet J, Mittelsten Scheid O, Paszkowski J. 2010. MOM1 and Pol-IV/V interactions regulate the intensity and specificity of transcriptional gene silencing. Embo j 29: 340–351.

Zhao H, Zhang W, Chen L, Wang L, Marand AP, Wu Y, Jiang J. 2018. Proliferation of Regulatory DNA Elements Derived from Transposable Elements in the Maize Genome. Plant Physiol 176: 2789–2803.

