## Supplementary material for "Natural diversity of heat-induced transcription of retrotransposons in *Arabidopsis thaliana*": FigureS

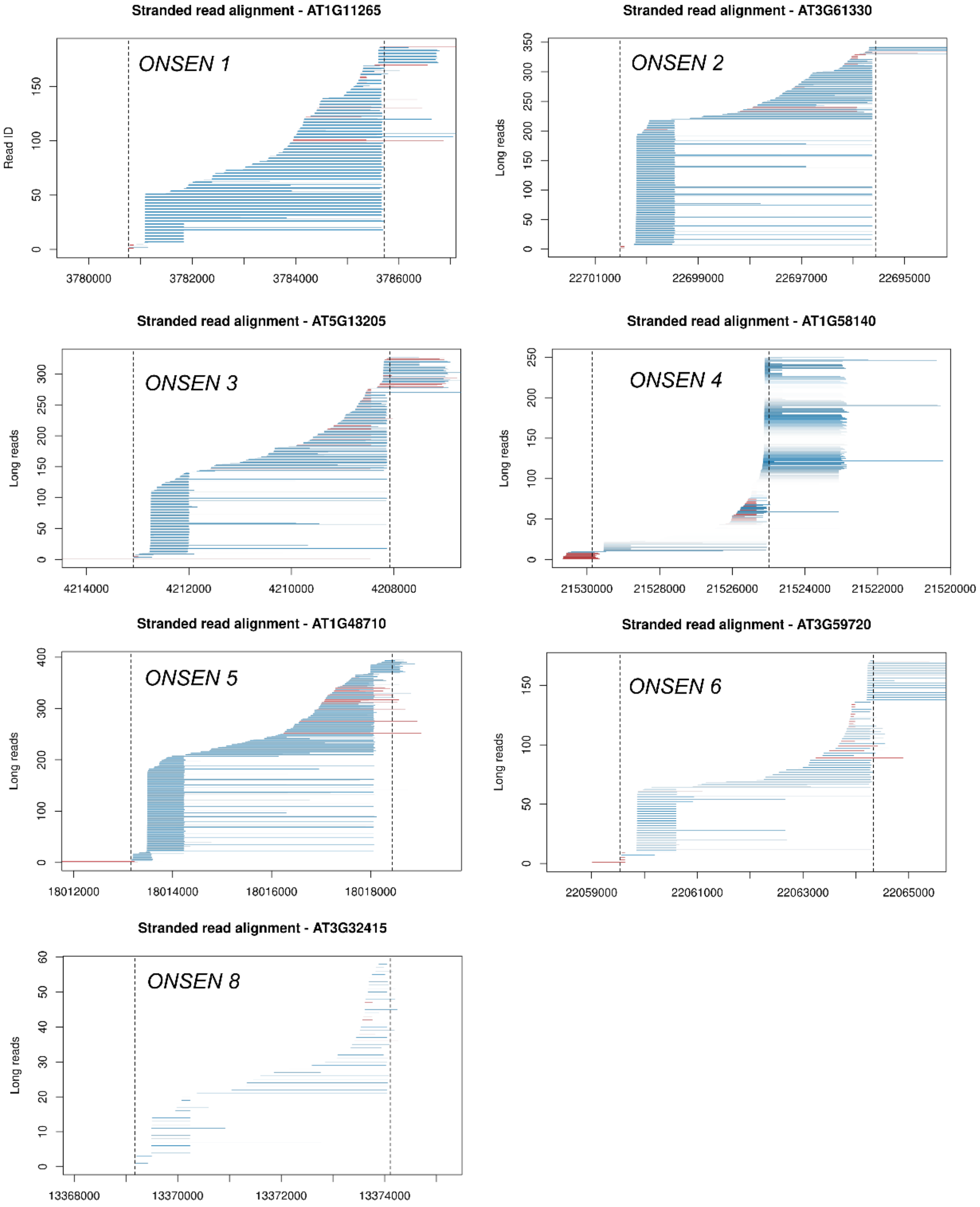


**Figure S1. Transcriptional profile of *ONSEN* copies in Col-0**

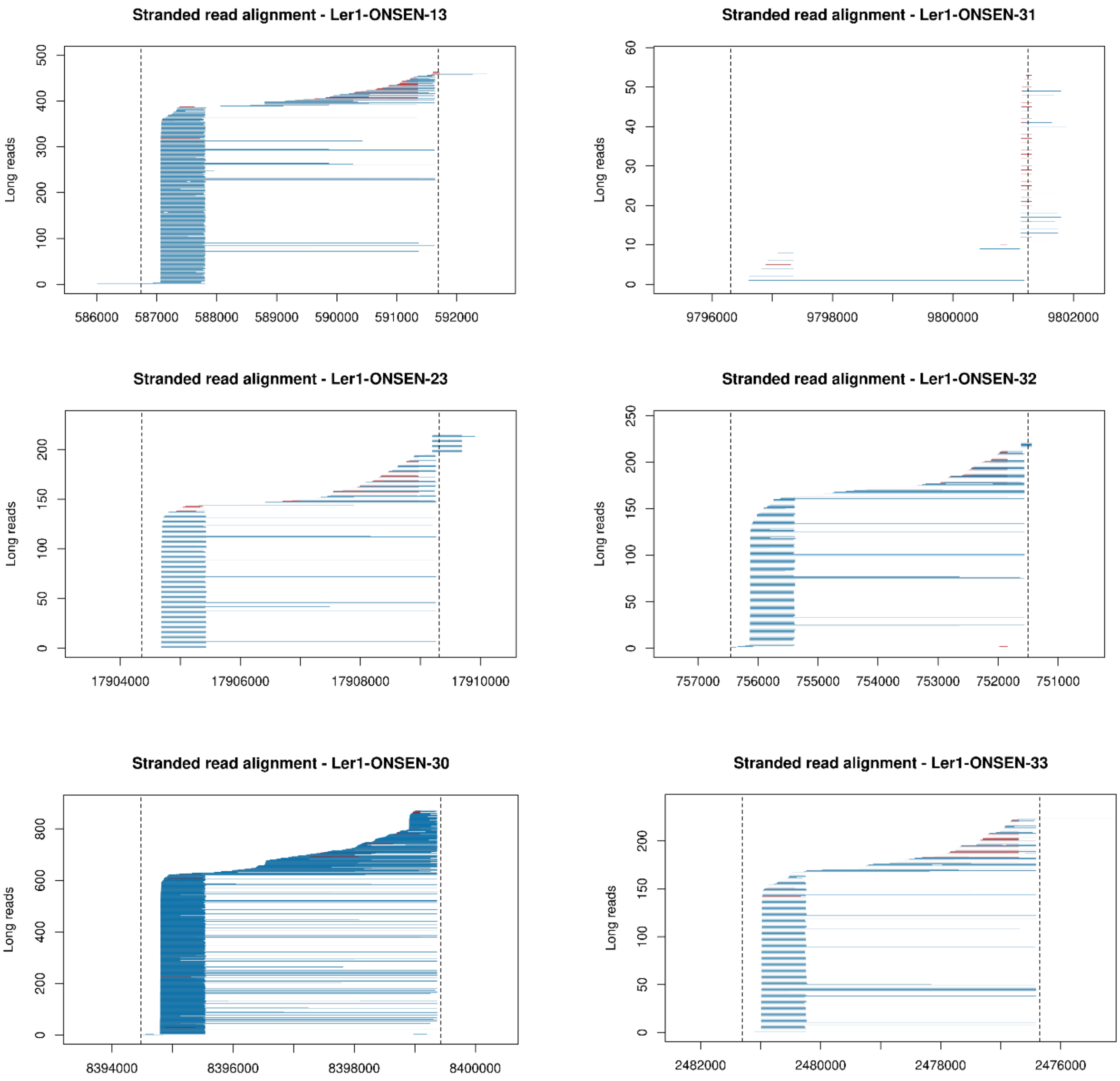


**Figure S2. Transcriptional profile of *ONSEN* copies in Ler-1**

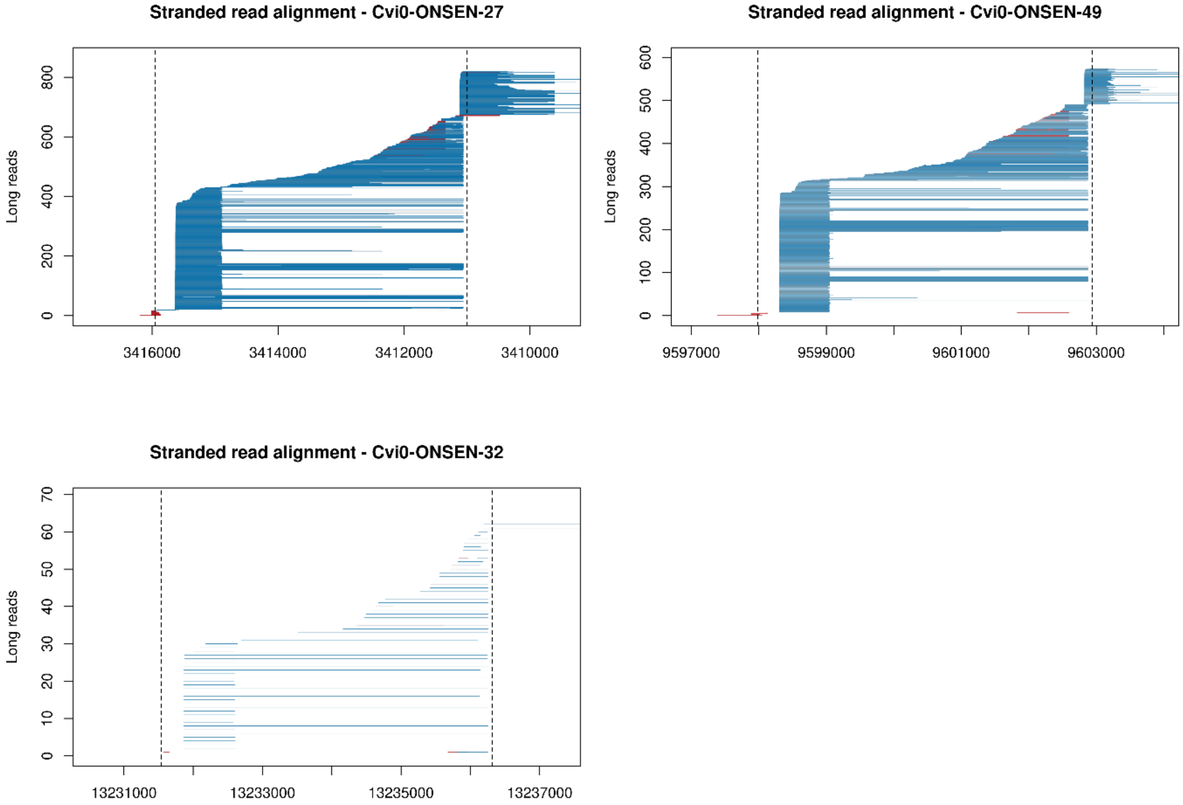


**Figure S3. Transcriptional profile of *ONSEN* copies in Cvi-0**

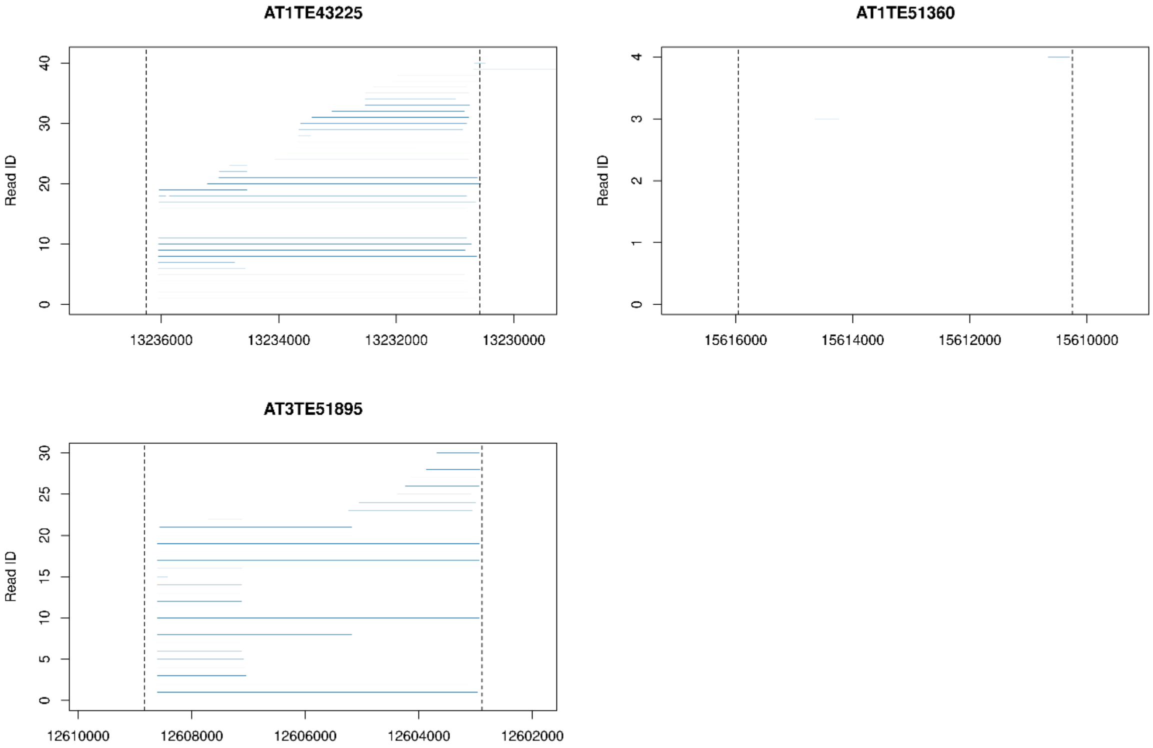


**Figure S4. Transcriptional profile of *Copia-35* copies in Col-0**

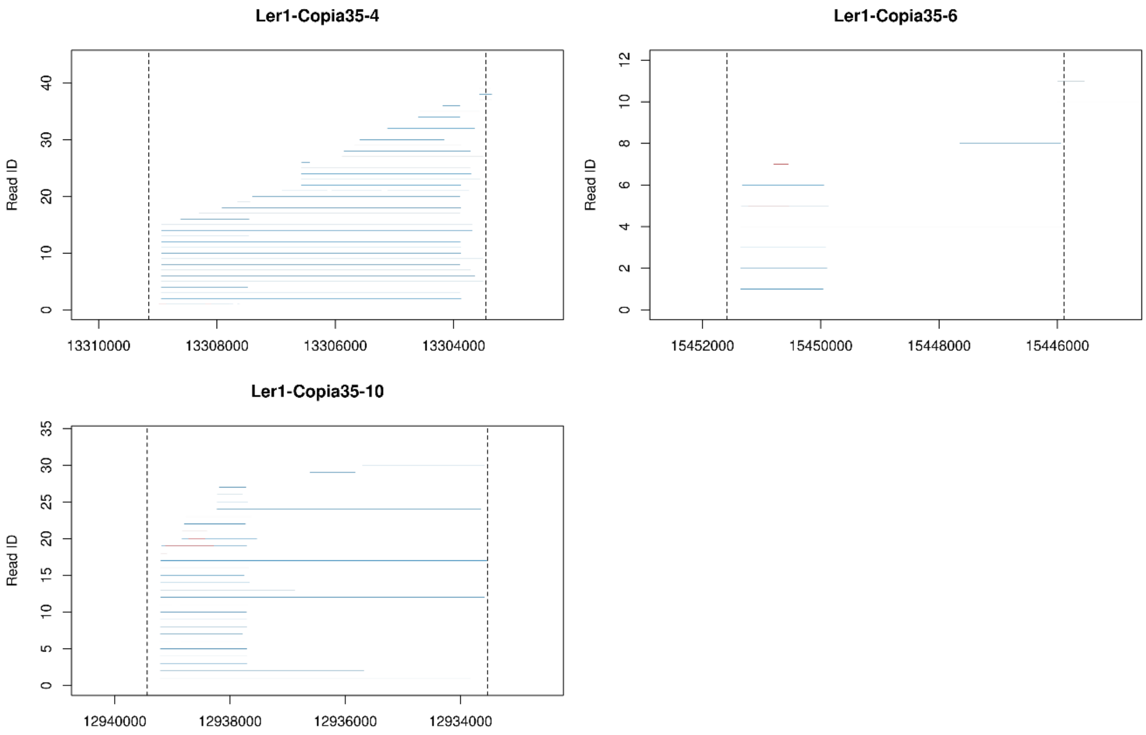


**Figure S5. Transcriptional profile of *Copia-35* copies in Ler-1**

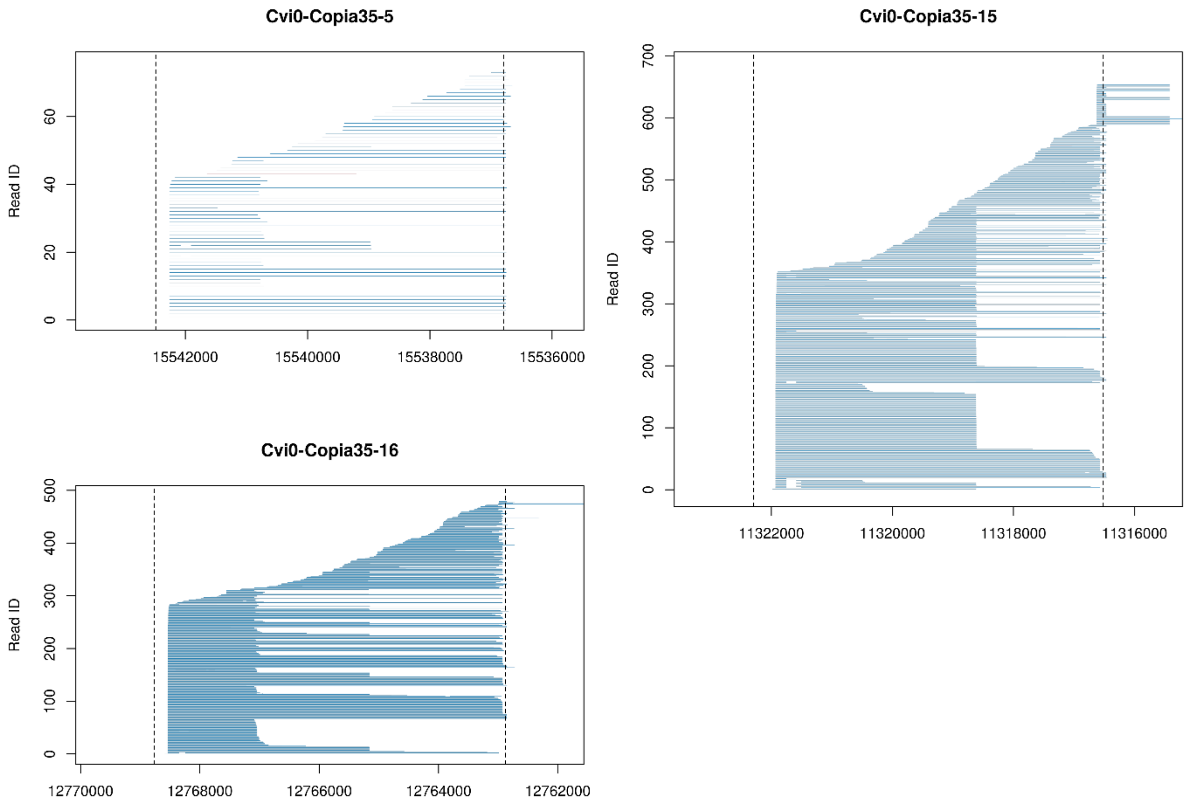


**Figure S6. Transcriptional profile of *Copia-35* copies in Cvi-0**

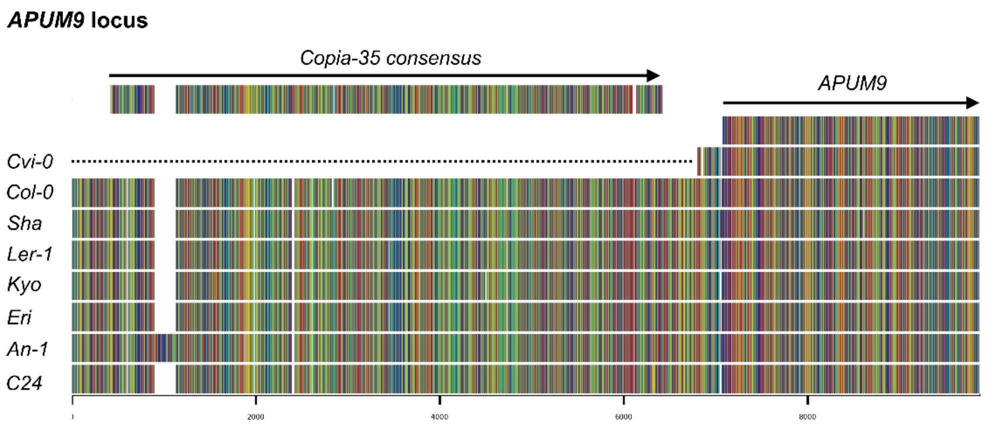


**Figure S7. *APUM9* locus of 8 PacBio assemblies.**

### Table S1. The RNA-Seq and ONT data quality (Uploaded separately)

### Table S2. Full-length *ONSEN* elements and their S1 Strength

| *ID* | Chrom | Start | End | S1 Reads | S2 Reads | S1 Strength S1/(S1+S2)*100% | S1/S2 | 3' Gene | 3' Gene reached by S2 reads |
|---|---|---|---|---|---|---|---|---|---|
| ***AT1G11265/ONSEN1*** | Chr1 | 3780765 | 3785721 | 138 | 19 | 87.90% | 7.26 | AT1G11280 | FALSE |
| ***AT1G21945/ONSEN7*** | Chr1 | 7717255 | 7722647 | NA | NA | NA | NA | AT1G21950 | FALSE |
| ***AT1G48710/ONSEN5*** | Chr1 | 18013162 | 18018435 | 321 | 18 | 94.70% | 17.83 | AT1G48730 | TRUE |
| ***AT1G58140/ONSEN4*** | Chr1 | 21524995 | 21529851 | 60 | 148 | 28.80% | 0.41 | AT1G58130 | TRUE |
| ***AT3G32415/ONSEN8*** | Chr3 | 13369174 | 13374108 | NA | NA | NA | NA | AT3G32425 | FALSE |
| ***AT3G59720/ONSEN6*** | Chr3 | 22059535 | 22064329 | 102 | 41 | 71.30% | 2.49 | AT3G59740 | FALSE |
| ***AT3G61330/ONSEN2*** | Chr3 | 22695566 | 22700522 | 291 | 11 | 96.40% | 26.45 | AT3G61310 | FLASE |
| ***AT5G13205/ONSEN3*** | Chr5 | 4208083 | 4213084 | 230 | 39 | 85.50% | 5.9 | AT5G13210 | FLASE |
| ***Ler1-ONSEN-13*** | Chr2 | 586740 | 591695 | 397 | 18 | 95.70% | 22.06 | ATLER-2G11780 | FLASE |
| ***Ler1-ONSEN-23*** | Chr3 | 17904360 | 17909316 | 146 | 22 | 86.90% | 6.64 | ATLER-3G66970 | TRUE |
| ***Ler1-ONSEN-30*** | Chr4 | 8394474 | 8399430 | 699 | 0 | 100% | inf | ATLER-4G38650 | FLASE |
| ***Ler1-ONSEN-31*** | Chr4 | 9796312 | 9801243 | NA | NA | NA | NA | ATLER-4G42665 | FLASE |
| ***Ler1-ONSEN-32*** | Chr5 | 751500 | 756456 | 179 | 8 | 95.70% | 22.38 | ATLER-5G12430 | TRUE |
| ***Ler1-ONSEN-33*** | Chr5 | 2476353 | 2481309 | 173 | 1 | 99.40% | 173 | ATLER-5G17680 | FLASE |
| ***Cvi0-ONSEN-27*** | Chr3 | 3410998 | 3415955 | 640 | 141 | 81.90% | 4.54 | ATCVI-3G20890 | TRUE |
| ***Cvi0-ONSEN-32*** | Chr3 | 13231539 | 13236316 | 51 | 3 | 94.40% | 17 | ATCVI-3G50870 | TRUE |
| ***Cvi0-ONSEN-49*** | Chr4 | 9597983 | 9602941 | 473 | 76 | 86.20% | 6.22 | ATCVI-4G40910 | TRUE |

### Table S3. Full-length *Copia-35* elements and their S1 strength

| *ID* | Chrom | Start | End | S1 Reads | S2 Reads | S1 Strength S1/(S1+S2)*100% | S1/S2 | 3' Gene | 3' Gene reached by S2 reads |
|---|---|---|---|---|---|---|---|---|---|
| *AT1TE43225* | Chr1 | 13230575 | 13236255 | 39 | 2 | 95% | 19.5 | AT1G35730 | FLASE |
| *AT1TE51360* | Chr1 | 15610250 | 15615952 | 3 | 1 | 75% | 3 | AT1G41830 | FLASE |
| *AT3TE51895* | Chr3 | 12602886 | 12608833 | 28 | 0 | 100% | 28 | AT3G30842 | FLASE |
| *Ler1-Copia35-4* | Chr1 | 13303453 | 13309151 | 42 | 2 | 95% | 21 | ATLER-1G48170 | FLASE |
| *Ler1-Copia35-6* | Chr1 | 15445886 | 15451583 | 10 | 2 | 83% | 5 | ATLER-1G55480 | FLASE |
| *Ler1-Copia35-10* | Chr3 | 12933534 | 12939434 | 34 | 0 | 100% | inf | ATLER-3G49220 | FLASE |
| *Cvi0-Copia35-5* | Chr1 | 15536790 | 15542484 | 76 | 0 | 100% | inf | ATCVI-1G55690 | FLASE |
| *Cvi0-Copia35-15* | Chr3 | 11316521 | 11322293 | 610 | 66 | 76% | 3.24 | ATCVI-3G43580 | TRUE |
| *Cvi0-Copia35-16* | Chr3 | 12762880 | 12768764 | 483 | 6 | 99% | 80.5 | ATCVI-3G48960 | TRUE |
